# Inhibition of selective autophagy by members of the herpesvirus ubiquitin-deconjugase family

**DOI:** 10.1101/2021.03.30.437649

**Authors:** Päivi Ylä-Anttila, Maria G. Masucci

**Affiliations:** Department of Cell and Molecular Biology, Karolinska Institutet, S-17165 Stockholm, Sweden

**Keywords:** autophagy, deubiquitinase, herpesviruses, large tegument protein, SQSTM1/p62

## Abstract

Autophagy is an important component of the innate immune response that restricts infection by different types of pathogens. Viruses have developed multiple strategies to avoid autophagy to complete their replication cycle and promote spreading to new hosts. Here we report that the ubiquitin deconjugases encoded in the N-terminal domain of the large tegument proteins of Epstein-Barr virus (EBV), Kaposi Sarcoma herpesvirus (KSHV) and human cytomegalovirus (HCMV), but not herpes simplex virus-1 (HSV-1), regulate selective autophagy by inhibiting the activity of the autophagy receptor SQSTM1/p62. We found that all the homologs bind to and deubiquitinate SQSTM1/p62 but with variable efficiency, which correlates with their capacity to prevent the colocalization of LC3 with SQSTM1/p62 aggregates and promote the accumulation of a model autophagy substrate. The findings highlight important differences in the strategies by which herpesviruses interfere with selective autophagy.

## Introduction

Autophagy is a protein degradation and recycling machinery that regulates cellular homeostasis and participates in the host defense against infection by capturing and destroying invading microorganisms (1, 2). Autophagy entails the sequestration of a portion of the cytoplasm in a double membraned vacuole which, upon fusion with a lysosome, causes the degradation of its content (1). The process can be either non-selective when a random portion of the cytoplasm is degraded, such as during starvation (3), or selective, when directed to cargoes that are marked for degradation by tags, such as ubiquitin or galectins, that are recognized by specific autophagy receptors (4). Known autophagy receptors include optineurin (OPTN) that recognizes ubiquitin coated protein aggregates, bacteria and mitochondria (5), calcium binding and coiled-coil domain 2/nuclear dot protein of 52kDa (CALCOCO2/NDP52) that recognizes galectin 8 on bacteria (6), sequestosome 1 (SQSTM1/p62) that recognizes a variety of ubiquitinated cargoes such as protein aggregates, lipid droplets, lysosomes, midbody rings, mitochondria, peroxisomes, zymogen granules and pathogens (7, 8, 9, 10, 11, 12, 13, 14), BCL2 and adenovirus E1B 19-kDa-interacting protein 3-BNIP3 like/NIP-3-like protein X (BNIP3-BNIP3L/NIX) that are mitochondrial membrane resident proteins functioning in mitophagy (15) and neighbor of BRCA1 gene 1 protein (NBR1) that recognizes ubiquitin on protein aggregates or peroxisomes (16, 17). A common feature of the autophagy receptors is their capacity to interact with the microtubule associated protein 1 light chain 3 (LC3) via a binding module known as LC3-interacting region (LIR), which promotes the wrapping of the growing autophagosomal membrane around the cargo (18).

Although generally considered as a protective cellular defense mechanism, autophagy can have pro-infection properties since many pathogens have evolved ways to modulate the process to their advantage. The strategies by which pathogens capture autophagy are beginning to be elucidated at the molecular level, which provides new insight on pathogenesis and highlights potential targets for therapeutic intervention (19). We have previously reported that Epstein-Barr virus ubiquitin-specific protease (USP or deubiquitinating enzyme, DUB) encoded in the N-terminal domain of the large tegument protein BPLF1 regulates autophagy (20). We found that BPLF1 binds to and deubiquitinates SQSTM1/p62, which hampers the recruitment of LC3 to SQSTM1/p62 positive structures. The deubiquitination of SQSTM1/p62 by BPLF1 is associated with accumulation of aggregation-prone mutant huntingtin (HTT) containing extended polyglutamine repeats (HTT-polyQ) and with failure to clear preformed aggregates, supporting the conclusion that BPLF1 can inhibit selective autophagy. A critical role for deubiquitination of the Lys420 residue located in the C-terminal domain of SQSTM1/p62 that mediates the interaction with both cargo and LC3 was revealed by the finding that the inhibitory effect could be rescued by co-expression of a mutant SQSTM1/p62 that mimics the active conformation induced by ubiquitination.

Although many of the BPLF1 substrates and interacting partners identified to date are shared by the BPLF1 homologs encoded by other herpesviruses, important differences have been observed. For example, subtle differences in the amino acid composition and charge of the otherwise relatively well conserved helix-2 were shown to prevent the interaction of the HSV-1 encoded homolog, UL36, with the 14-3-3 molecular scaffolds, resulting in failure to modulate the activity of the ubiquitin ligase TRIM25 (21). Here we set out to investigate whether the ubiquitin deconjugases encoded by EBV, HCMV, KSHV and HSV-1 share the capacity to regulate selective autophagy by interfering with the activity of the SQSTM1/p62 receptor. We found that similar to BPLF1, the catalytic domains of HCMV-UL48, KSHV-ORF64 and HSV-1-UL36 bind to and deubiquitinate SQSTM1/p62 but the efficiency and ubiquitin chain specificity of the viral DUBs was remarkably different. Furthermore, the HSV-1 encoded UL36 failed to inhibit the recruitment of LC3 to SQSTM1/p62 positive structures and did not promote the accumulation of mutant HTTpolyQ into insoluble aggregates. Thus, it appears that the different lifestyles of herpesviruses may have promoted the development of distinct strategies for interfering with selective autophagy.

## Materials and methods

### Chemicals

DL-dithiothreitol (DTT, D0632), N-ethylmaleimide (NEM, E1271), iodoacetamide (I1149), IGEPAL CA-630 (NP40, I3021), Triton X-100 (T9284), bovine serum albumin (BSA, A7906), sodium dodecyl sulfate (SDS, L3771), Tween-20 (P9416), ethylenediaminetetraacetic acid disodium salt dehydrate (EDTA, E4884), Trizma base (Tris, 93349) and ciprofloxacin (I7850) were purchased from Sigma-Aldrich. Complete protease inhibitor cocktail (04693116001) and phosphatase inhibitor cocktail (04906837001) were purchased from Roche Diagnostic.

### Antibodies

Antibodies and their manufacturers were: mouse anti-FLAG clone M2 (WB 1:4000, IF 1:400; F1804) and rabbit anti-LC3B (WB 1:1000, IF 1:200; L7543) were from Sigma-Aldrich; goat anti-FLAG (1:400; ab1257), rabbit anti-SQSTM1/p62 (IP 1-2 μg/mg of lysate; ab101266) and rabbit anti-GFP (WB 1:2000, ab290) were from Abcam; mouse anti-SQSTM1/p62 (WB 1:1000 IF 1:200; 610832) was from BD Biosciences; rabbit anti-SQSTM1/p62 (WB 1:1000, 8025S) was from Cell Signaling Technology; mouse anti-HA.11 clone 16B12 (WB 1:1000, 901501) was from Biolegend; mouse anti-HA clone 12CA5 (WB 1:2000; 11583816001) from Roche; mouse IgG isotype control (14-4732-82) was from Invitrogen; Alexa Fluor 488-, 555-, 594- and 647-conjugated secondary antibodies were from Thermo Fisher (A21206, A31570, A11032 and A21447, respectively).

### Plasmids

Plasmids encoding 3xFLAG-BPLF1 (amino acid residues 1-235), the catalytic mutant BPLF1-C61A, KSHV-ORF64 (amino acids 1-265) and HA-tagged ubiquitin were described previously (22), (23). Plasmid encoding codon optimized 3xFLAG-tagged HSV-UL36 (amino acid residues 1-293) was synthesized at Gene Universal Inc. DE, USA and 3xFLAG-HCMV-UL48 was kindly provided by Luka Cicin-Sain, Helmholtz Center for Infection Research, Braunschweig, Germany (24). Plasmids pRK5-HA-UbK48 and pRK5-HA-UbK63 that encode for ubiquitin polypeptides where all Lys residues are mutated to Arg except for Lys48 or Lys63 were kindly provided by Harald Wodrich, Laboratoire de Microbiologie Fondamentale et Pathogenicite UMR-CNRS, University of Bordeaux. HTTQ109-GFP plasmid was a gift from Nico Dantuma, Department of Cell and Molecular Biology, Karolinska Institutet. The coding sequence of the SQSTM1/p62-K420R,E409A mutant was excised from the plasmid pET28a-p62-K420R/E409A-Flag (25) kindly provided by Ronggui Hu (Institute of Biochemistry and Cell Biology, Shanghai Institutes for Biological Sciences, China) and inserted between HindIII and NotI restriction sites of the HA-SQSTM1/p62 expression vector (26) (Addgene, 28027; deposited by Qing Zhong). The HA-SQSTM1/p62-K7A mutant was described previously (20).

### Cell lines and transfection

HeLa cells (ATCC RR-B51S) cells were cultured in Dulbecco’s modified Eagle’s medium (DMEM, Sigma-Aldrich, D6429), supplemented with 10% FCS (Gibco-Invitrogen, 10270-106), and 10 μg/ml ciprofloxacin, and propagated at 37°C in a 5% CO_2_ incubator. Plasmid transfection was performed using the JetPEI (Polyplus transfection, 101-40N) or Lipofectamine 2000 kits (Life technologies, 11668019) as recommended by the manufacturer.

### Immunoblotting and immunoprecipitation

For immunoblotting and co-immunoprecipitation cells harvested 48 h post transfection were lysed in NP40 lysis buffer (50 mM Tris-HCl pH 7.6, 150 mM NaCl, 5mM MgCl2, 1 mM EDTA, 1 mM DTT, 1% Igepal, 10% glycerol) supplemented with protease and phosphatase inhibitors from Roche (04693116001, 4906837001) and deubiquitinase inhibitors (20 mM NEM and 20 mM Iodoacetamide). Protein concentration was measured with a Lowry protein assay kit (Bio-Rad Laboratories). For co-immunoprecipitation, the cell lysates were incubated for 4 h with anti-FLAG agarose affinity gel (Sigma, A-2220). To compensate for expression differences, three-fold higher amounts of proteins were used for UL36 and ORF64 IP. Precipitated complexes were washed with lysis buffer and eluted with the FLAG peptide (Sigma, F4799) at a concentration of 400 μg/ml. For co-immunoprecipitation of endogenous SQSTM1/p62 the cell lysates were incubated for 2-4 h with specific antibody followed by 1-2 h with protein-G coupled Sepharose beads (GE Healthcare, 17-0885-01). To resolve protein complexes for denaturing immunoprecipitation, cell pellets were lysed in 100 μl NP-40 lysis buffer supplemented with 1% sodium dodecyl sulfate (SDS) for 30 min at 4°C followed by addition of NP-40 buffer to reach a final concentration of 0.1% SDS. Immunocomplexes were washed with lysis buffer and elution was performed by boiling for 5 min in 2x SDS-PAGE loading buffer. Equal amounts of proteins were fractionated in a polyacrylamide Bis-Tris 4– 12% gradient gels (Invitrogen, NP0321PK2). After transfer to poly-vinylidene difluoride (PVDF) membranes (Millipore), the blots were blocked in Tris-buffered saline containing 5% non-fat milk and 0.1% Tween-20 and incubated with primary antibodies either for 1 h at room temperature or overnight at 4°C followed by incubation for 1 h with the appropriate horseradish peroxidase-conjugated secondary antibodies. The complexes were visualized by chemiluminescence (ECL; Pierce, 32106).

### Filter-trap assay

Cells were harvested 48 h after transfection, resuspended in PBS (Biowest, X0515-500) containing protease inhibitor cocktail and samples were snap frozen and stored at −20°C. Lysates were sonicated with a QSonica Q125 sonicator with settings 20% amplitude, pulsating 1s on/1s off total time of 30 sec. After sonication protein amounts were measured with Bio-Rad Lowry kit according to the manufacturer’s instructions. Samples with equal protein concentrations were prepared by dilution in PBS containing protease inhibitors and SDS was added to a final concentration of 1%. The lysates were suctioned through a cellulose acetate membrane (GE health care, 10404180) using a Bio-Rad Bio-Dot apparatus and immunoblotting was performed as described. Images were analyzed with the ImageJ software.

### Immunofluorescence and confocal microscopy

HeLa cells were grown to semi-confluence on glass cover slips in Dulbecco’s modified Eagle’s medium (Sigma, D6429) containing 10% fetal calf serum and 10 μg/ml ciprofloxacin and transfected with the indicated plasmids using the JetPEI or Lipofectamine 2000 kit as recommended by the manufacturers. After treatment for the indicated time the cells were fixed in 4% paraformaldehyde (Merck, 100496,) and permeabilized with cold methanol at −20°C for 2 min or 0.1% TX-100 in PBS for 5 min at room temperature followed by blocking with 0.12% glycine (Fisher Scientific, G46-1) in PBS for 10 min. Blocking in 3% bovine serum albumin (BSA, Sigma, A7906) in PBS for 15 min at room temperature was performed before antibody labeling. The cells were labeled in 3% BSA-PBS using rabbit anti-LC3, mouse anti-SQSTM1/p62 and goat anti-FLAG or mouse anti-FLAG antibodies followed by the appropriate Alexa Fluor 488, 555, 594 or 647 conjugated secondary antibodies. The nuclei were stained with 2.5 μg/ml DAPI (Sigma, D9542) in PBS for 10 min and the coverslips were mounted cell side down on object glasses with Mowiol (Calbiochem, 475904) containing 50 mg/ml 1,4-diazabicyclo[2.2.2]octane (Dabco; Sigma, D-2522) as anti-fading agent. The slides were imaged using a confocal scanning laser microscope (Zeiss LSM880) and 1 μm optical sections were acquired.

### Image analysis

Images were analyzed with the Fiji software. The size of SQSTM1/p62 structures and colocalization of LC3 and SQSTM1/p62 labels was analyzed by creating a mask based on thresholding the SQSTM1/p62 fluorescence. The number and size of the SQSTM1/p62 structures were counted with the analyze-particles function and LC3 fluorescence was measured through the mask by acquiring the mean gray value for each structure. Empty-vector transfected sample was used as a control. Changes in the level of LC3 colocalization with SQSTM1/p62 were assessed by comparing the mean gray values in test-transfected cells with the mean grey values of all structures detected in empty-vector transfected cells.

## Results

### The herpesvirus DUBs interact with SQSTM1/p62 and regulate its ubiquitination

We have previously reported that the catalytic domain of the EBV large tegument protein BPLF1 inhibits selective autophagy (20). BPLF1 deubiquitinated SQSTM1/p62 and this was associated with impaired recruitment of LC3 to SQSTM1/p62 positive structures and inability to clear cytosolic protein aggregates. In order to investigate whether the BPLF1 homologs encoded by other herpesviruses share the capacity to interfere with selective autophagy via interaction with SQSTM1/p62, Hela cells were transfected with FLAG-tagged versions of the catalytic domains of EBV-BPLF1, HSV-UL36, HCMV-UL48 and KSHV-ORF64 and cell lysates were immunoprecipitated with anti-FLAG coated beads. The precipitates were fractionated on polyacrylamide gels and blots were probed with antibodies specific for the FLAG tag and SQSTM1/p62. In spite of significantly different levels of expression, all viral homologs reproducibly showed robust interaction with SQSTM1/p62 (Figure 1A). Of note, due to the poor expression of UL36 and ORF64 higher amounts of cell lysates were used for the input, hence the stronger SQSTM1/p62 bands in the blots shown in Figure 1A. We then asked if the homologs also share the capacity to deubiquitinate SQSTM1/p62. To this end, and to explore whether different types of polyubiquitin chains may be preferentially recognized, the viral enzymes were co-transfected in HeLa cells together with plasmids expressing ubiquitin mutants where all Lys residues except Lys48 or Lys63 were substituted with Arg (HA-UbK48 and HA-UbK63). The transfected cells were lysed in denaturing conditions to disrupt protein-protein interactions, and endogenous SQSTM1/p62 was then immunoprecipitated with a specific antibody. The total amounts of polyubiquitinated proteins and ubiquitinated SQSTM1/p62 were visualized by probing blots of the input and immunoprecipitated samples with antibody against the HA-tag (Figures 1B and 1D). As previously reported, overexpression of the viral enzymes was associated with a global decrease in the levels of polyubiquitinated proteins (22), which affected to similar extent Lys48- and Lys63-linked chains (Figures 1B and 1D, input). However, consistent differences were observed in the capacity of the viral enzymes to remove polyubiquitin chains from SQSTM1/p62 (Figures 1B and 1D, IP). While BPLF1, UL48 and ORF64 acted on both types of chains with comparable efficiency, UL36 preferentially removed Lys48-linked chains and exhibited much weaker activity against Lys63-linked chains (Figures 1C and 1E).

**Figure 1.**
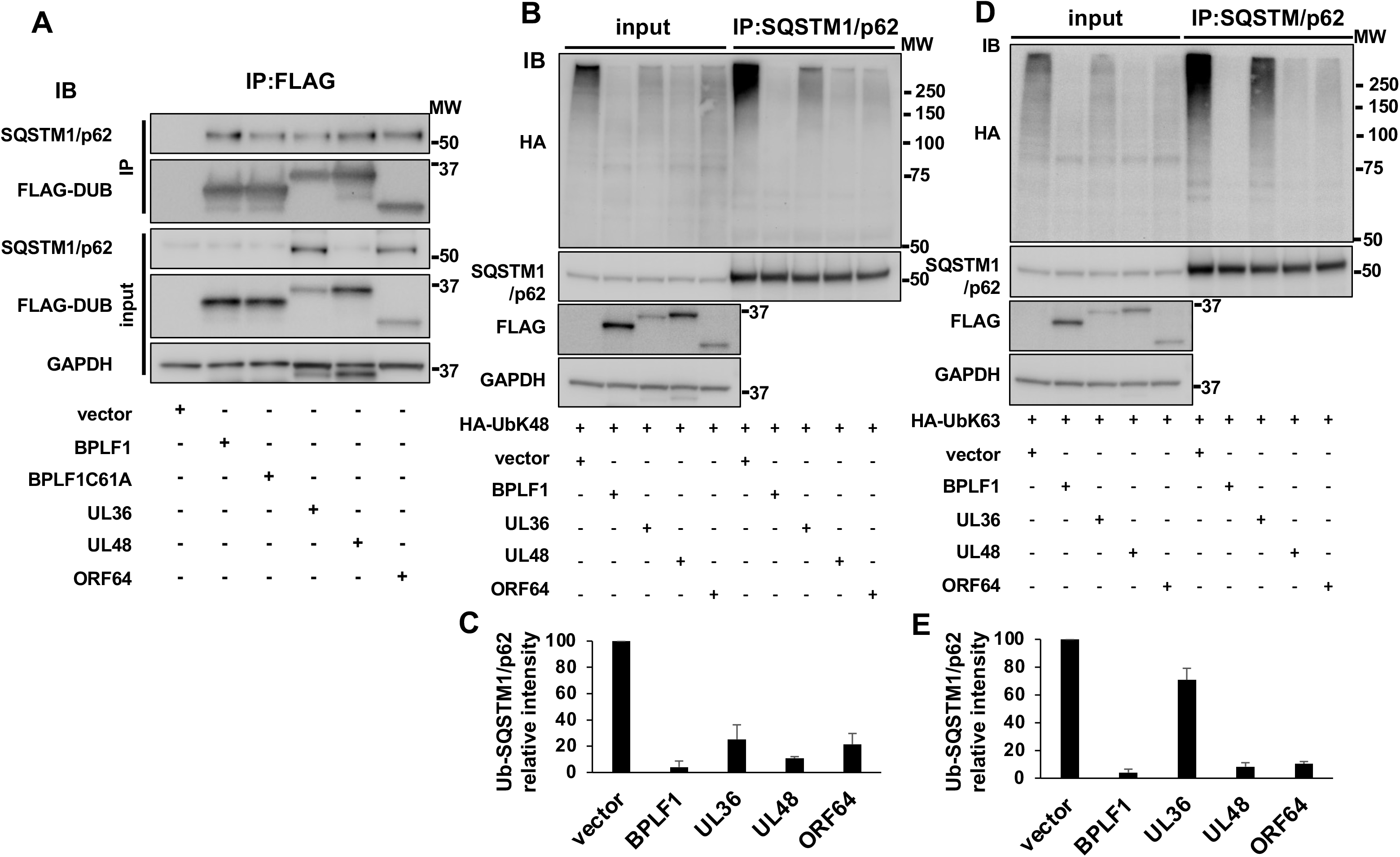
EBV-BPLF1, HSV-1-UL36, HCMV-UL48 and KSHV-ORF64 interact with SQSTM1/p62 but differentially affect its ubiquitination. (A) HeLa cells were transfected with FLAG-tagged BPLF1, UL36, UL48 or ORF64 and cell lysates were immunoprecipitated with anti-FLAG coated beads. Comparable levels of co-immunoprecipitated SQSTM1/p62 were detected in all samples. Western blots from one representative experiment out of two where the viral proteins were tested in parallel are shown. (B, D) BPLF1, UL36, UL48 or ORF64 were co-transfected in HeLa cells together with HA-UbK48 or HA-UbK63 and endogenous SQSTM1/p62 was immunoprecipitated with specific antibody in denaturing conditions. Polyubiquitin chains were detected by probing western blots with antibodies to the HA tag. Representative western blots from one representative experiment out of two are shown. (C, E) The intensity of the ubiquitin smears was quantified by densitometry in two independent experiments. The Mean ± SD of smear intensity in test samples relative to the vector control are shown. The viral enzymes removed K48-linked polyubiquitin chains from SQSTM1/p62 with comparable efficiency (B, C), whereas UL36 exhibited significantly weaker activity against K63-linked chains (D, E).

### Differences in SQSTM1/p62 deubiquitination correlate with LC3 colocalization

The capacity of SQSTM1/p62 to deliver polyubiquitinated cargo to the autophagosome via interaction with LC3 is essential for selective autophagy (27). To investigate whether HCMV-UL48, KSHV-ORF64 and HSV1-UL36 share with BPLF1 the capacity to prevent the colocalization of SQSTM1/p62 with LC3, HeLa cells transfected with plasmids expressing the viral enzymes were co-stained with antibodies specific for LC3 and SQSTM1/p62. As previously reported (20), the expression of catalytically active BPLF1 was associated with a marked decrease in the size of SQSTM1/p62 structures relative to vector transfected cells, whereas the catalytic mutant BPLF1-C61A where the active Cys residue was substituted with Ala had no effect. A decrease of dot size was observed also in cells expressing HCMV-UL48 and, to a minor extent KSHV-ORF64 but not in cells expressing HSV1-UL36 (Figure 2A, 2B). In line with previous observation that the small SQSTM1/p62 dots fail to colocalize with LC3 (20), a significant reduction in the number of SQSTM1/p62 aggregates decorated with LC3 was observed in cells expressing BPLF1 and HCMV-UL48, whereas HSV1-UL36 and KSHV-ORF64 had no appreciable effect (Figure 2A, 2C).

**Figure 2.**
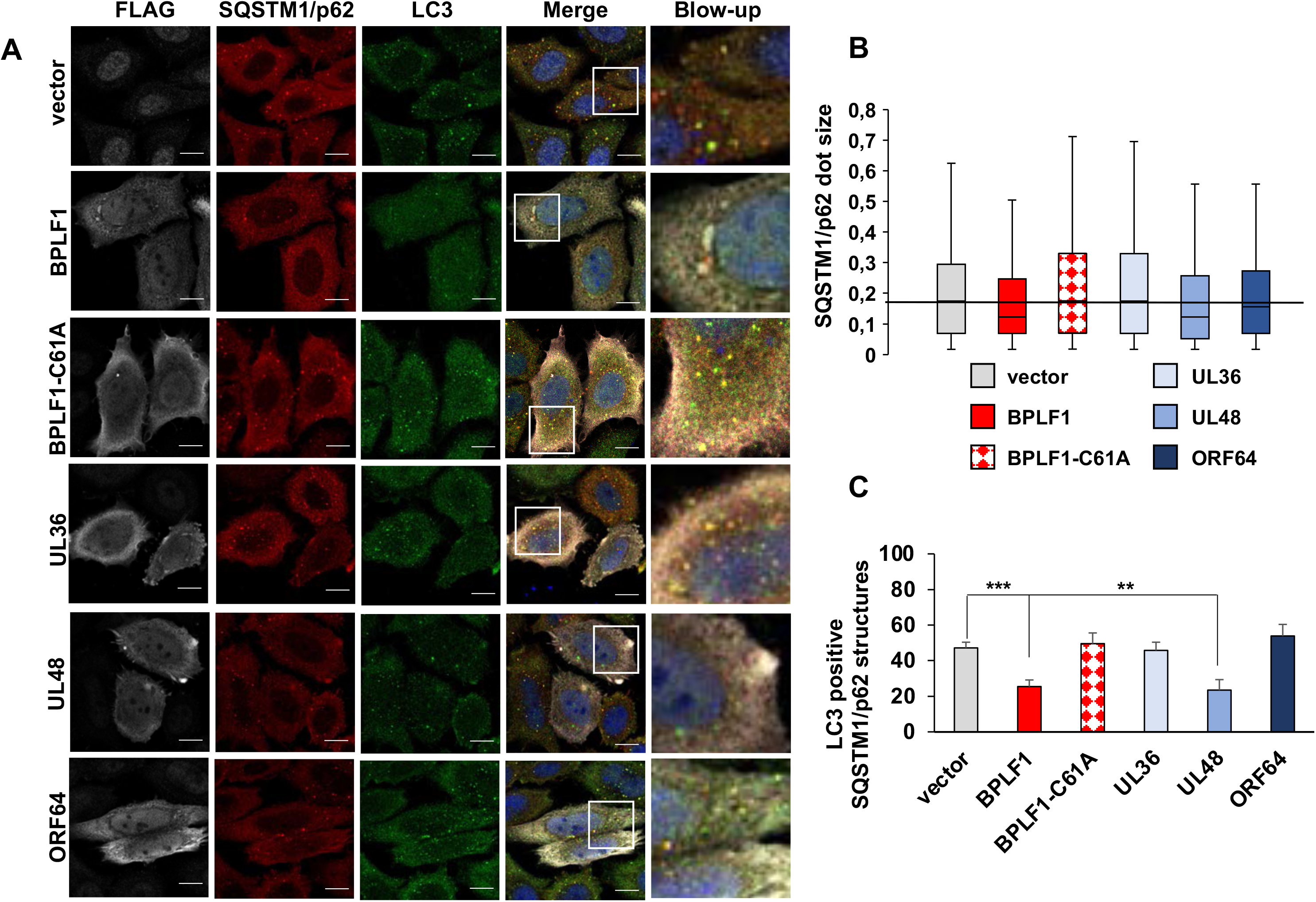
EBV-BPLF1 and HCMV-UL48 inhibit the recruitment of LC3 to SQSTM1/p62 positive structures. HeLa cells were transfected with FLAG-tagged BPLF1, UL36, UL48 or ORF64 and endogenous LC3 and SQSTM1/p62 were detected by immunofluorescence. (A) representative macrographs illustrating the colocalization of SQSTM1/p62 (red) with LC3 (green) in cells expressing the viral enzymes (gray). The nuclei were stained with DAPI (blue). Size bars = 10 μm. (B) The size of SQSTM1/p62 structures were notably smaller in the cells expressing BPLF1, UL48 and ORF64 compared to empty vector expressing control cells and cells expressing either BPLF1-C61A or UL36. (C) The colocalization of LC3 with SQSTM1/p62 dots was measured using the ImageJ software. The mean ± SD of the % LC3 positive SQSTM1/p62 positive dots in two independent experiments where a minimum of 34 cells were analyzed is shown.

### The viral DUBs differentially affect the autophagy-mediated clearance of SQSTM1/p62 client proteins

The activity of SQSTM1/p62 in selective autophagy is regulated by the ubiquitination of Lys residues located in the C-terminal and N-terminal domains that control the interaction with ubiquitinated cargo (25) and the formation of oligomeric filaments that facilitate the sequestration of cargo into LC3-coated membranes (28), respectively. In order to investigate whether the different effect of the viral DUBs on SQSTM1/p62 ubiquitination may impact selective autophagy we monitored their effect on the autophagy-mediated clearance of a model substrate. An aggregation-prone GFP-tagged huntingtin mutant containing a 109 amino acid long poly-glutamine repeat (HTT109Q-GFP) was co-transfected in HeLa cells together with plasmid expressing FLAG-tagged BPLF1, UL36, UL48 and ORF64, and the accumulation of HTTQ109-GFP aggregates was monitored by filter-trap assays. Cell lysates prepared 48 h after transfection were blotted onto cellulose acetate membranes that are permeable to SDS soluble proteins, using a dot-blot apparatus, and trapped aggregates were detected by probing the membranes with antibodies to GFP (Figure 3A, 3B). As previously reported (20), expression of BPLF1 caused the accumulation of HTTQ109-GFP aggregates. To account for the differences in expression levels, increasing amounts of plasmids encoding the viral enzymes were transfected together with a constant amount of the HTTQ109-GFP plasmid. Expression of EBV-BPLF1, HCMV-UL48 and KSHV-ORF64 was accompanied by a dose-dependent accumulation of HTTQ109 aggregates. The effect of KSHV-ORF64 was somewhat weaker, which is in line with the failure to significantly impair the recruitment of LC3 to SQSTM1/p62 positive structures. In contrast, HTTQ109-GFP aggregates did not accumulate in cells expressing HSV1-UL36 even upon strong overexpression of the viral enzyme (Figure 3C), suggesting that this viral DUB does not inhibit SQSTM1/p62-dependent autophagy.

**Figure 3.**
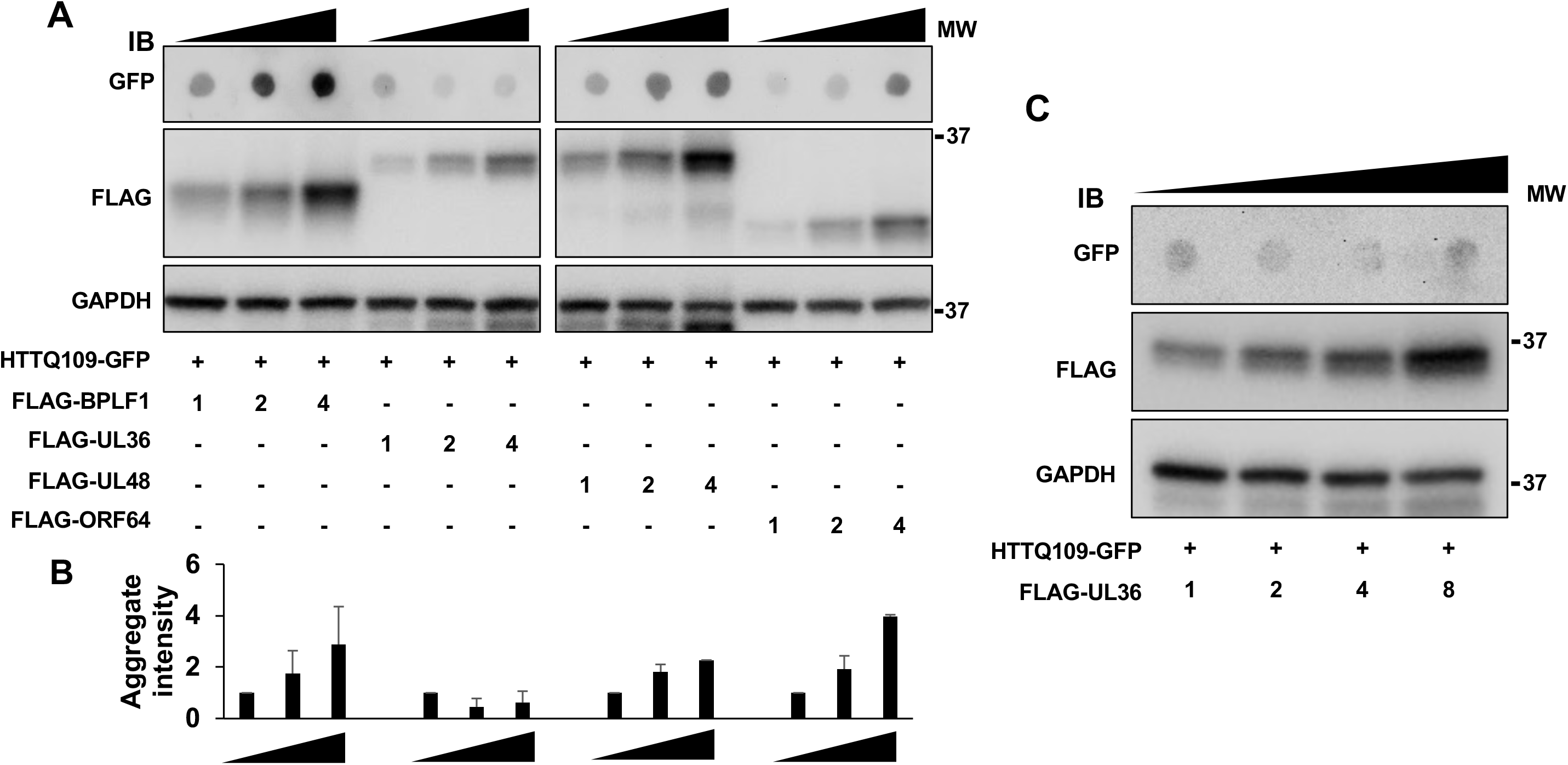
Effect of the viral DUBs on the autophagy-mediated clearance of SQSTM1/p62 client proteins. HeLa cells were co-transfected with plasmids expressing aggregation-prone HTTQ109-GFP and increasing amounts of the viral DUBs. The accumulation of HTTQ109-GFP aggregates was monitored 48 h after transfection by filter-trap assays followed by probing of the blotted membranes with a GFP-specific antibody. (A) A dose-dependent accumulation of HTTQ109-GFP aggregates was observed in cells expressing EBV-BPLF1, HCMV-UL48 and KSHV-ORF64 while HSV1-UL36 had no appreciable effect. The numbers correspond to the amoints of transfected plasmids in μg. Representative blots from one representative experiment out of two where the DUB were tested in parallel are shown. (B) The intensities of the GFP dots were quantified using the ImageJ software and mean ± SD of dot intensity in two independent experiments is shown. (C) HSV1-UL36 failed to induce the accumulation of HTTQ109-GFP aggregates even at the highest concentrations tested.

Ubiquitination of SQSTM1/p62 at Lys420 was shown to prevent electrostatic interactions between the Lys420 and Glu409, which promotes a conformational change in C-terminal domain that allows loading of the ubiquitinated cargo (25). We have previously shown that overexpression of a SQSTM1/p62 mutant where the Lys and Glu residues are substituted with Arg and Ala, respectively (SQSTM1/p62-E409A, K420R) can override the capacity of BPLF1 to inhibit selective autophagy, suggesting that deubiquitination of SQSTM1/p62 Lys420 may be critical for the effect of the viral enzyme (20). To investigate whether this residue is also targeted by the homologs that share with BPLF1 the capacity to inhibit selective autophagy, HeLa cells transfected with plasmids expressing HTTQ109-GFP and either HCMV-UL48 or KSHV-ORF64 were co-transfected with increasing amounts of the SQSTM1/p62-E409A, K420R mutant (Figure 4). The SQSTM1/p62-E409A, K420R mutant rescued in a dose-dependent manner the capacity of HCMV-UL48 to promote the accumulation of HTTQ109-GFP aggregates, whereas a SQSTM1/p62-K7A mutant that is still dependent on K420 ubiquitination for cargo loading and is oligomerization deficient lacked this capacity, confirming that the rescue is not an artifact of overexpression (Figure 4A and 4B). The weaker inhibitory activity of KSHV-ORF64 required high amount of transfected plasmid to achieve detectable levels of HTTQ109-GFP aggregates, which did not allow testing the effect of a large excess of the SQSTM1/p62-E409A, K420R mutant. In spite of this technical limitation, clearance of the HTTQ109-GFP aggregates was rescued even by relatively low amounts of the SQSTM1/p62-E409A, K420R mutant (Figure 4C).

**Figure 4.**
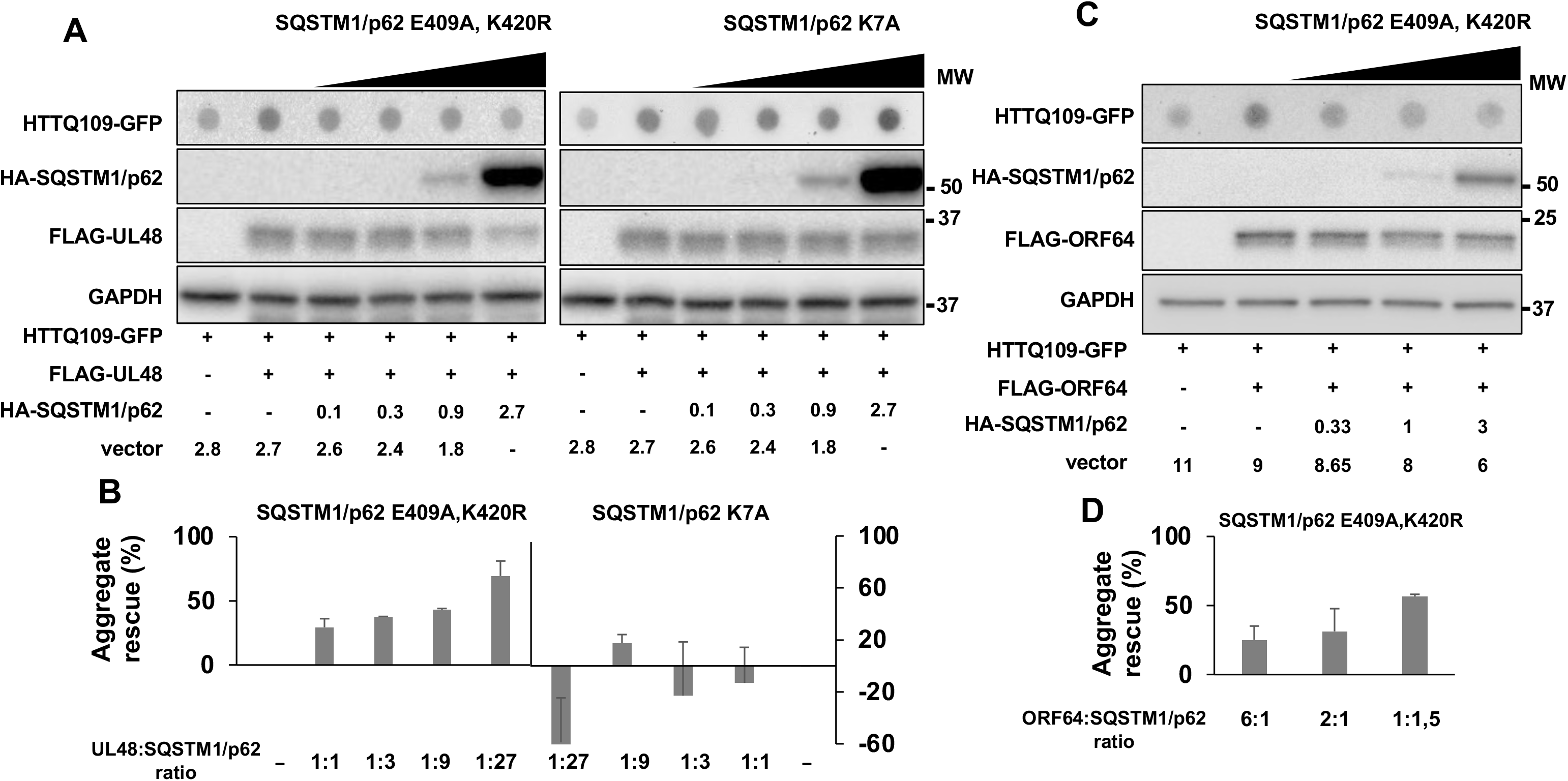
The accumulation of HTTQ109-GFP aggregates is rescued by a constitutively active SQSTM1/p62 mutant. HeLa cells were co-transfected with fixed amounts of plasmids encoding the viral DUBs and HTT109-GFP and increasing amounts of plasmids encoding the indicated SQSTM1/p62 mutants. The accumulation of HTTQ109-GFP aggregates was assessed by filter-trap assays 48 h after transfection. (A) The accumulation of HTTQ109-GFP aggregates induced by HCMV-UL48 was rescued by expression of the SQSTM1/p2-E409A, K420R mutant but not by the SQSTM1/p62-K7 mutant. The numbers indicated the amount of transfected plasmids in μg. Blots from one representative experiment out of two are shown. (B) Quantification of dot intensity recorded in two independent experiments. (C) The SQSTM1/p2-E409A, K420R mutant rescued the accumulation of HTTQ109-GFP aggregates induced by KSHV-ORF64. Blots from one representative experiment out of two are shown. (D) Quantification of dot intensity recorded in two independent experiments.

## Discussion

While compelling evidence points to a key role of selective autophagy in the clearance of both the incoming viruses and newly synthesized viral proteins (19), different viruses have evolved distinct strategies for counteracting this cellular defense to promote their replication and spread. In this study, we report on important differences in the capacity of human herpesviruses to regulate SQSTM1/p62-dependent selective autophagy by interfering with SQSTM1/p62 ubiquitination. We found that ubiquitin deconjugases encoded in the N-terminal domain of the large tegument proteins of HSV-1, HCMV, EBV and KSHV interact with and deubiquitinate SQSTM1/p62. However, while this resulted in inhibition of the SQSTMI/p62-dependent clearance of a model substrate in cells expressing the enzymes encoded by HCMV, EBV, and KSHV, the HSV-1 encoded enzyme had no appreciable effect. The failure to inhibit selective autophagy correlates with the inability of the HSV-1 enzyme to remove Lys63-linked polyubiquitin chains from SQSTM1/p62 and to prevent the colocalization of SQSTM1/p62 aggregates with LC3 decorated autophagic membranes. These findings highlight a previously unrecognized contribution of Ly63-linked polyubiquitination in the regulation of SQSTM1/p62 activity that is differentially exploited by herpesviruses to counteract selective autophagy.

We have previously shown that SQSTM1/p62 binds to and is a substrate of the EBV encoded enzyme BPLF1 (20). Here we found that this property is shared by the homologs encoded by other human herpesviruses, albeit with important differences in the type of ubiquitin chains that can be removed by the viral enzymes (Figure 1). Most notably, the HSV-1 encoded UL36 showed a significantly reduced capacity to remove Lys63-linked ubiquitin chains from SQSTM1/p62. Conflicting data have been reported on the efficiency and ubiquitin chain specificity of UL36 (29, 30). However, our findings are not explained by a general or selective failure of enzymatic activity since overexpression of UL36 induced comparable levels of global deubiquitination, with no apparent preference for the type of ubiquitin linkage. It is noteworthy that, although structurally similar, the catalytic domains of the herpesvirus tegument proteins show very little amino acid sequence conservation (29). The only relatively well-conserved domain is the solvent-exposed helix-2 that is involved in the interaction of BPLF1 with cullin ligases (23) and with the 14-3-3 molecular scaffolds (31). Interestingly, molecular modeling suggests that the helix-2 of UL36 is shorter compared to the corresponding domain in BPLF1, UL48 and ORF64, and the solvent exposed residues are primarily negatively charged, while the homologs share a domain organization with negatively charged N-terminus and positively charged C-terminus (21). It is tempting to speculate that, while directly responsible for the failure of UL36 to interact with 14-3-3 (21), these subtle structural differences may also modulate the interaction of UL36 with putative substrates, which may also dictate the recognition of different types of ubiquitin chains. A detailed mapping of the interaction site(s) of SQSTM1/p62 with the herpesvirus homologs would be required to clarify this issue.

SQSTM1/p62 functions in the autophagic degradation of ubiquitinated cargo by physically linking the cargo to the autophagic machinery via binding to autophagosome-anchored LC3 (32). The interaction with the cargo is mediated by the C-terminal ubiquitin associated domain (UBA) and is dependent on the ubiquitination of critical residues, which prevents SQSTM1/p62 homodimerization and the consequent occlusion of the cargo-binding site (25). We have previously reported that the deubiquitination of SQSTM1/p62 by BPLF1 is associated with the accumulation of small SQSTM1/p62 structures that do not co-localize with LC3, and with failure to clear cytosolic aggregates of mutant HTTQ109 (20). Clearance of the HTTQ109 aggregates was rescued by overexpression of the mutant SQSTM1/p62-E409A, K420R, pointing to a scenario where deubiquitination of K420 by BPLF1 hampers the capacity of SQSTM1/p62 to load ubiquitinated cargo and to promote its sequestration into LC3-decorated autophagosomes. We found that these properties of BPLF1 are shared by the homologs encoded by the HCMV and KSHV, but not by the HSV-1 encoded UL36 (Figure 2, 3, and 4). Of note, the weaker effect of KSHV-ORF64 on the size of SQSTM1/p62 aggregates and LC3 colocalization (Figure 2) is likely to be the result of comparably poor expression since accumulation of HTTQ109 aggregates could be achieved by transfecting higher amounts of ORF64 plasmid (Figure 3A, 3B). In contrast, even the highest amounts of overexpressed UL36 failed to inhibit the clearance of HTTQ109 aggregates (Figure 3), pointing to a true difference in the capacity of this viral DUB to modulate selective autophagy.

Our findings have several interesting implications. First, the behavior of the viral enzymes highlights an important role of different types of ubiquitin chain in the regulation of SQSTM1/p62 activity. Several ligases were shown to attach polyubiquitin chains on SQSTM1/p62 with different outcomes (33, 34, 35). In particular, opposite effects were observed upon polyubiquitination of Lys7 in the C-terminal PB1 domain of SQSTM1/p62; while the TRIM21-mediated attachment of Lys63-linked polyubiquitin chains was shown to inhibit the oligomerization of SQSTM1/p62 and the sequestration of cargo (33), attachment of the same type of chains by NEDD4 induced the formation of large SQSTM1/p62-structure and promoted autophagy (34). Thus, the same modification may have different effects depending on the cellular context. Conceivably, the failure of UL36 to inhibit the formation of Lys63 linked chains on Lys7 could explain the retained formation of large SQSTM1/p62 aggregates that colocalize with LC3 in UL36 expressing cells, but differences in the ubiquitination status of this residue are not sufficient to explain the concomitant failure to inhibit selective autophagy.

The capacity of the SQSTM1/p62-E409A, K420R mutant to rescue the inhibitory effects of BPLF1, UL48, and ORF64 suggests that deubiquitination of Lys420 is critical for the activity of viral enzymes. Several ubiquitin ligases, including the Keap1/Cullin3 ligase (35), and under ubiquitin-stress conditions the UBE2D2/UBE2D3 conjugating enzymes (25), were shown to polyubiquitinate the Lys420 residue, but the type of ubiquitin chains was not investigated. It is noteworthy that Keap1/Cullin3 can form Lys48-, Lys63- and even mixed Lys48-Lys63-linked ubiquitin chains on different substrates (36, 37). As illustrated by the cartoon shown in Figure 5, our findings point to a scenario where different types of chains may be formed on SQSTM1/p62-K420, which could have distinct effects on the fate and function of the protein. Lys48-linked, Lys63-linked or mixed chains will all prevent the formation of heterodimers mediated by electrostatic interaction between E409 and K420 with consequent activation of the receptor function. Thus, elimination of all types of chains may be required to stabilize the formation of homodimers that prevent cargo binding and inhibit autophagy; whereas persistent Lys63-linked ubiquitination appears to allow autophagy. Interestingly, the free UBA domain shows strong preference for Lys63-ubiquitinated cargo (38). Hence, intermolecular competition of Lys63-ubiquitinated SQSTM1/p62 for cargo binding could inhibit autophagy, while intramolecular blockade of the UBA receptor by long Lys-63-linked chains may redirect SQSTM1/p62 to other functions such as the regulation of apoptosis (39), oxidative stress (40), or NF-kB dependent immune responses (41). Although not within the scope of this study, our findings emphasize the importance of a thorough dissection of the types of ubiquitin chains attached to SQSTM1/p62 in order to better understand their contribution to the regulation of the autophagy receptor.

**Figure 5.**
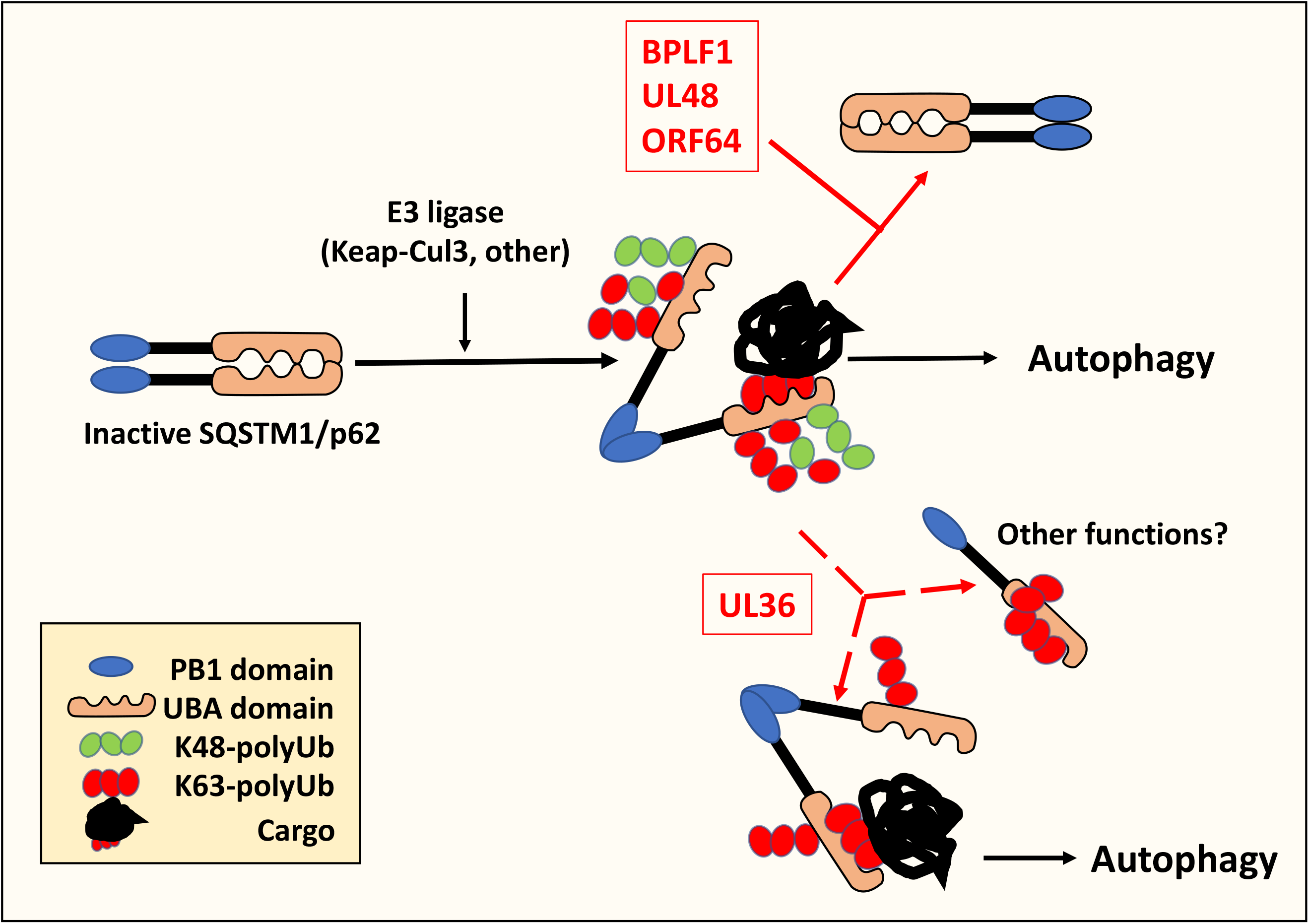
Model of SQSTM1/p62 regulation by the herpesvirus encoded DUBs. Cellular ubiquitin ligases attach different types of polyubiquitin chains to the autophagy receptor SQSTM1/p62. Ubiquitination of Lys420 inhibits the formation of SQSTM1/p62 homodimers by preventing electrostatic interaction with the Glu409 residue on adjacent SQSTM1/p62 molecules. This promotes an open conformation that allows the recruitment of ubiquitinated cargo to the UBA domain. The deubiquitinating enzymes encoded in the N-terminus of HSV-1-UL36, HCMV-UL48, EBV-BPLF1 and KSHV-ORF64 bind to and deubiquitinate SQSTM1/p62. UL48, BPLF1 and ORF64 remove both Lys48 and Lys63-linked ubiquitin chains locking SQSTM1/p62 in heterodimers that cannot bind ubiquitinated cargo, which results in failure to recruit the autophagic machinery and inhibits autophagy. In UL36 expressing cells, the persistence of Lys63-linked chains is sufficient to prevent the formation of heterodimers and activates SQSTM1/p62-dependent autophagy. In addition, intramolecular blockade of the UBA by Lys63-linked chains may redirect SQSTM1/p62 to other functions.

In spite of their common ancestry and shared capacity to establish both latent and productive infections, herpesviruses differ significantly in their target spectrum, which is likely to require different strategies for manipulation of the host cell environment. Alpha-herpesviruses, such as HSV-1, replicate very efficiently in epithelial cells and establish latency in non-replicating neurons whereas beta- and gamma-herpesviruses can both replicate and establish latency in hematopoietic cells and their replication cycle is significantly slower (42). We have previously shown that the HSV-1 encoded UL36 differs from the homologs encoded by EBV, HCMV and KSHV in its failure to inhibit innate immune responses by counteracting the activity of the TRIM25 ubiquitin ligase (21). Here we have described a second property of UL36 that sets it apart from the homologs. This finding further emphasizes the presence of substantial differences in the strategies adopted by herpesviruses for manipulating the host cell environment and counteracting the host antiviral defense.

## Acknowledgements

We are grateful to Drs. Harald Wodrich, Nico Dantuma, Luka Cicin-Sain and Ronggui Hu for providing plasmids and technical advice.

## Data availability

All data that support the findings of this study are contained within the manuscript and are available on reasonable request.

## Abbreviations

BPLF1: BamH1 fragment left open reading frame-1
EBV: Epstein-Barr virus
GFP: green fluorescent protein
HTT: Huntingtin
MAP1LC3/LC3: microtubule associated protein 1 light chain 3
PB1: Phox and Bem1 domain
PE: phosphatidylethanolamine
SQSTM1/p62: sequestosome 1
UBA: ubiquitin associated domain

